# Epidermal TRPV4 ion channels regulate UVB induced sunburn by triggering inflammmasome activation and ERK signaling

**DOI:** 10.1101/2021.01.09.426056

**Authors:** Carlene Moore, Shinbe Choi, Gene Moon, Jennifer Zhang, Yong Chen, Wolfgang Liedtke

## Abstract

Skin inflammation is an evolutionary-honed protective mechanism that serves to clear noxious cues and irritants and initiate regeneration. Calcium-permeable transient-receptor-potential (TRP) ion channels have critical functions in sensory transduction which is sensitized in skin inflammation. Skin sensory transduction relies on skin-innervating sensory neurons in the dorsal root ganglion (DRG), but also on innervated keratinocytes (KC). The multimodally-activated TRPV4 is robustly expressed in KC, where it can readily be activated by Ultraviolet-B (UVB). Our goal was to deconstruct keratinocyte TRPV4-mediated signaling, specifically how TRPV4 can facilitate inflammatory injury, thus lowering pain thresholds and rendering KC into pain-generator cells. We wanted to uncover the effect of TRPV4-mediated signaling on UVB-induced inflammasome activation in KC given the powerful impact of the activated inflammasome on pro-inflammatory/pro-algesic secretory signaling, using mouse models and cultured human KC. In mice, our evidence suggests that TRPV4 functions as calcium-permeable channel upstream of the KC inflammasome. Furthermore, we found that UVB induced activation of TRPV4 caused rapid - within minutes - **E**xtracellular Signal **R**egulated **K**inase (ERK) phosphorylation, caspase-1 activation and Interleukin-1ß (IL-1ß) secretion. In human primary KC we demonstrated that UVB induced secretion of IL-1ß was dependent on the NLR family pyrin domain containing 1 (NLRP1) inflammasome. Direct chemical TRPV4 activation could also activate NLRP1 and to lesser extent NLPR3. Building on our previous work, we now define at increased resolution TPRV4-dependent forefront signaling mechanisms in KC in response to UVB, showing TRPV4 upstream of the NLRP1 inflammasome, subsequent rapid ERK activation and pro-inflammatory/pro-algesic secretory function.

## INTRODUCTION

The skin, the largest vertebrate organ, provides critical barrier protection of a consistent *milieu interne* and defends against potentially harsh external environments (1–4). Keratinocytes (KCs) of the epidermal surface epithelium not only provide a structural barrier but function as the organismal forefront to interact with and sense the environment (2). This forefront signaling can activate/modulate the immune system and skin innervating sensory neurons, the latter function perhaps underappreciated. The skin surface epithelium is closely associated with the peripheral axons of the sensory neurons of the dorsal root ganglia and trigeminal ganglia (DRG, TG), that are endowed with the full sensory transduction capacity including itch and pain (2–5). Amidst a backdrop of suggestive findings (2,6–8), we wondered by which mechanism epidermal KCs sensitize pain transduction in response to naturally occurring irritating cues. To elucidate these mechanisms, we took advantage of a mouse sunburn model which induces a state of lowered sensory thresholds evoked by UVB over-exposure (9–11). UVB-evoked lowering of sensory thresholds shares major hallmarks of pathological pain for a self-limited period of several days.

TRPV4 is a widely expressed, multimodally-activated non-selective cation channel expressed in both innervated epithelia and sensory neurons, such as epidermal KCs and skin-innervating DRG/TG sensory neurons. It has also been found relevant for pain transduction and transmission and experimental itch (12–19). TRPV4 responds to osmotic, mechanical, injurious and inflammatory cues (20), also to UVB, and it functions in inflammation, nerve damage and UVB-induced sensitization (21,22). Previously we showed that following UVB over-exposure, mice with genetically-engineered *Trpv4* deletions in skin KCs, with inducibility of the *Trpv4* knockdown, were less sensitive to painful thermal and mechanical stimuli vs control mice (23). Mechanistically, we found that epidermal TRPV4 can orchestrate UVB-evoked tissue damage through regulation of the expression of potent pro-inflammatory and pro-algesic mediators, namely IL-6 and endothelin-1. This finding clearly indicates that TRPV4-mediated Ca^++^-influx into epidermal KCs evokes secretion of pro-inflammatory factors, with the primary stress provided, UVB over-exposure, a known inflammasome activator. This begets the question whether TRPV4-mediated Ca^++^-influx activates the KC’s inflammasomes.

Inflammasomes are large multi-protein complexes that assemble in response to microbial, immune-mediated and other cellular injury, and trigger an inflammatory cascade culminating in caspase-1 activation and production of pro-inflammatory cytokines, e.g. IL-1ß and IL-18, with IL-1ß a proven pain-evoking mediator. Inflammasomes have been linked to various skin conditions including photo-aging, contact dermatitis, rosacea, atopic dermatitis and skin cancer, for all of which TRPV4 has been claimed to have some form of involvement (23–29). At a basic level, inflammasome activation has been shown to depend on Ca^++^-signaling (30,31) yet we do not know the molecular identity of the Ca^++^-entry machinery. This prompted us to investigate whether TRPV4 plays a significant role in inflammasome activation, how inflammasome activation depends on TRPV4, and in case this could be demonstrated, to identify relevant underlying mechanisms that result in release/secretion of pro-algesic and pro-inflammatory mediators by KC. We therefore conducted experiments to address this question, and, in case of affirmative results, to identify molecular mechanisms down-stream of TRPV4-mediated Ca^++^-influx, such as activation of MAP-kinases and identification of the isoform of the activated inflammasome that depends on TRPV4 (32)

## RESULTS

### Keratinocyte-specific ablation of *Trpv4* and topical inhibition of epidermal TRPV4 reduced keratinocyte IL-1ß expression in response to UVB in mice

First, we intended to test whether TRPV4 regulates the inflammasome by measuring established inflammatory effector molecule, caspase-1, and inflammatory cytokine IL-1ß. UVB irradiation of KCs has been shown to evoke inflammasome activation which activates the proteolytic effector, caspase-1, which then cleaves pro-IL-1ß to become active IL-1ß. We asked whether caspase-1 and IL-1ß expression were *Trpv4*-dependent (33). With UVB, we noted a significant upregulation of caspase-1 in the epidermis of control mice. In contrast, for *Trpv4* knockout animals, which refers to pan-null *Trpv4*^−/−^ and keratinocyte conditional-inducible *Trpv4*^−/−^ (KC-*Trpv4*(iKO)) mice, this regulation was absent (Fig.1 A-C). Interestingly, caspase-1 could not be detected via western blotting in *Trpv4*^−/−^ skin, indicating its expression was dependent on *Trpv4*. In agreement with these findings, we found that IL-1ß was not upregulated in the skin in response to UVB in *Trpv4*^−/−^ and KC-*Trpv4*(iKO)) mice (Fig.1 D-F). These findings indicate that in live mammals, *Trpv4* critically regulates expression and thus function of caspase-1 in skin, so that IL-1ß fails to be generated by KC in response to UVB. Furthermore, induced loss-of-function of *Trvp4* in KC also leads to failure of the skin to upregulate IL-1ß expression. Thus KC TRPV4 appears necessary for a basic inflammasome effector mechanism in-vivo.

**Figure 1.**
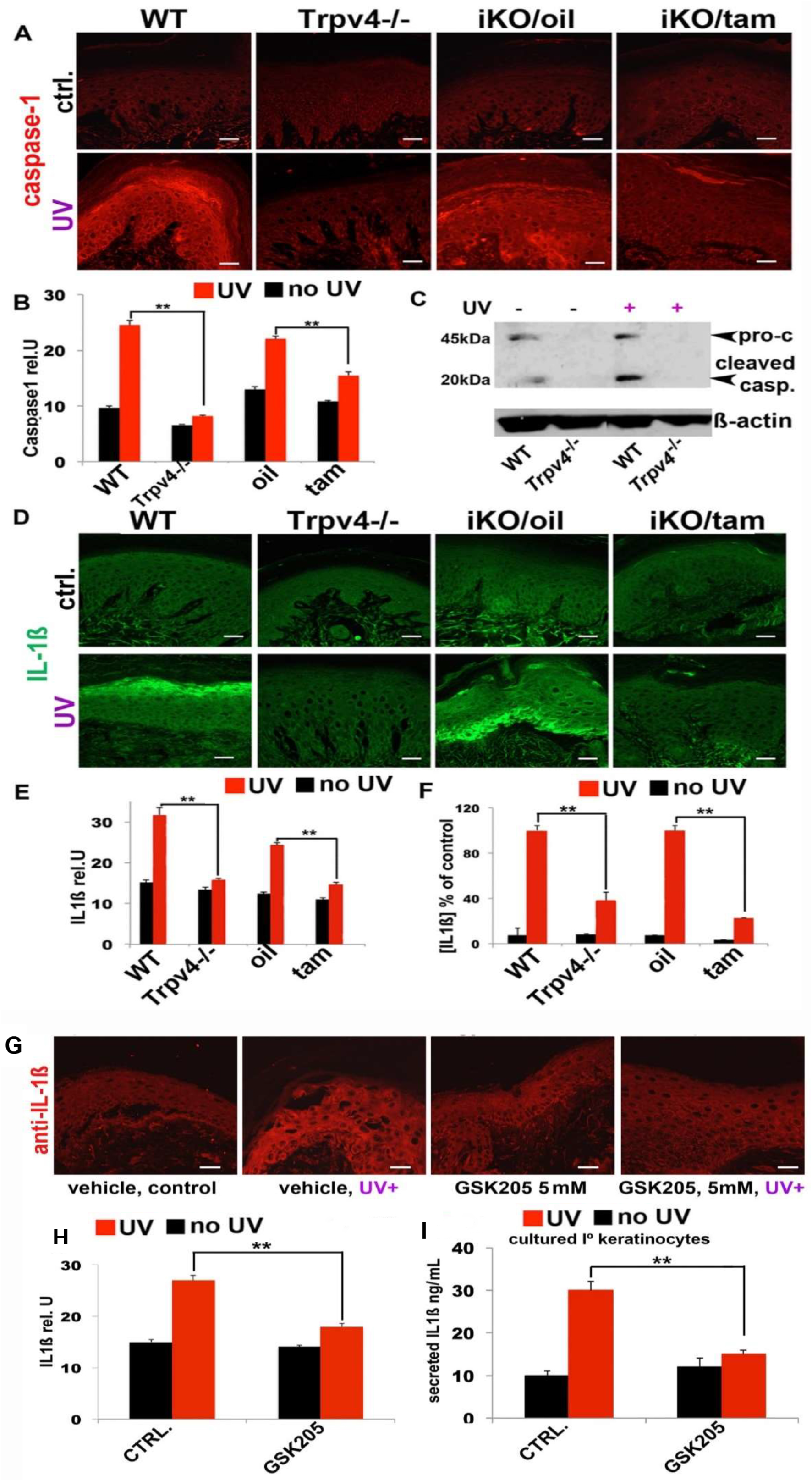
Keratinocyte-specific ablation of *Trpv4* and topical inhibition of epidermal TRPV4 reduced keratinocyte caspase1 and IL-1ß expression in response to UVB in mice. Immuno-histochemical analysis demonstrates UVB-mediated inflammasome activation in keratinocytes in mice depends on TRPV4. (A) Micrographs show caspase −1 immunolableing of representative skin footpads sections in response to UVB, sampled 48 hours after UVB treatment. Note that Caspase-1 is up-regulated in mouse KCs as a marker of inflammasome activation in both the WT and the oil treated iKO mice. Caspase 1 immunofluorescence reveals a reduced ability of TRPV4-deficient and the skin targeted tamoxifen treated iKO mice to elevate keratinocyte caspase1 in response to UVB exposure. (B) Quantifications of caspase-1 immunolableling. Densitometries are for *n* = 3 per group, showing significant up-regulation for WT and oil treated iKO, mice and lack thereof for *Trpv4^−/−^* and tam treated mice. The data are expressed as mean and SEM **p<0.01 *t* test. (Scale bar, 20μm) (C) Western blot of lysates from hindpaw skin showing an increase in caspase 1 cleavage in the skin of WT UVB treated skin however caspase1 expression was markedly reduced in *Trpv4^−/−^* mice and expression remained unchanged with UVB treatment. Note that caspase-1 levels, in particular cleaved caspase-1 (lower band), are elevated in UVB-exposed WT cells, but there is a complete absence of both procaspase-1 and cleaved caspase-1 in MPKCs from *Trpv4*^−/−^ mice. (D) IL1ß is up-regulated in mouse KCs as a marker of inflammasome activation. Anti-IL1ß immunofluorescence reveals a reduced ability of TRPV4-deficient mice to elevate keratinocyte IL1ß in response to UVB exposure similar to panel A. (Scale bar, 20μm). (E) Quantifications for IL-1ß immunolabeling. Densitometries are for *n* = 3 per group, showing significant up-regulation for WT and oil treated iKO, mice and lack thereof for *Trpv4^−/−^* and tam treated mice. **p<0.01 *t* test. (F) IL1ß ELISA concentrations in interstitial fluid derived from mice footpad of hind paw 48 hours after UV treatment. Note strong up-regulation in WT and oil-treated iKO mice after UVB, in contrast significant attenuation in *Trpv4*^−/−^ and tam-treated iKO mice. *n*=5 mice/group, ** p<0.01 *t* test. (G) External topical application of TRPV4 inhibitor GSK205 attenuates IL1ß expression. GSK205-treatment attenuates keratinocyte expression of IL-1ß in UVB-exposed footpad – representative micrographs of skin sections of UVB-exposed skin after UVB treatment. Bars=20μm. (H) GSK205-treatment attenuates keratinocyte expression of IL-1ß in UVB-exposed footpad - quantifications. Bar diagrams show densitometry results from *n*=3 mice/group, ** p<0.01 *t* test. (I) GSK205-treatment attenuates secretion of IL-1ß by UVB-exposed MPKCs. IL-1ß concentrations in supernatant (ELISA), are shown in response to UVB. Cells were cultured +/− 5μM GSK205. Note **p<0.01. Two-tailed *t* test was used for statistical analysis.

Loss-of-function is genetically-encoded in *Trpv4* null mice. To confirm that ion channel function of TRPV4 is the relevant mechanism for defective IL-1ß expression, we targeted TRPV4 by topical application of small molecule TRPV4 inhibitor GSK205. In our previous study we demonstrated unambiguously that topical GSK205, to paw skin, had no off-target effects in our UVB tissue injury/pain model, using pan-null *Trpv4*^−/−^ mice (23). With results shown in Fig. 1G-H, we topically applied GSK205 to mouse hind paw skin footpads and then exposed the animals to UVB. When the tissue was collected 48 hours after UVB treatment, the vehicle-treated mice showed an increase in epidermal IL-1ß expression, GSK205 topical treatment attenuated the increase in IL1ß expression, as observed in the *Trpv4* gene-targeted mice. In agreement with and in extension of these findings, GSK205 also blocked IL-1ß secretion from primary cultured mouse KCs.

### TRPV4 activation by UVB induces ERK phosphorylation, then triggers extracellular release of IL-1ß and TNFα in skin KCs

Our group has previously demonstrated that ERK signaling downstream of TRPV4 was critical for formalin-evoked irritant pain, histaminergic itch, and air pollution particulate matter-evoked airway irritation (34–36). To determine if ERK signaling was involved in the inflammatory response of UVB-absorbing skin KCs, we UVB-irradiated the back skin of WT mice. We found increased ERK phosphorylation in skin when assessed 15 minutes after UVB treatment, which was attenuated in *Trpv4*^−/−^ pan-null mice and KC-*Trpv4*(iKO) mice (Fig.2A, B). With a robust signal at the 15-minutes time-point, we sought to establish a higher-resolution time course of ERK phosphorylation in human derived primary KCs (HPKCs). In these cells we found that as early as 5 minutes after UVB treatment ERK was robustly phosphorylated, then decreased at the 15 minutes’ time point and was no longer detectable at the 60 minutes’ time point (Fig.2C). This upregulation was TRPV4 dependent as TRPV4-inhibitory compounds GSK205 and the more potent compound 16-8 (37) eliminated the increase in pERK. To further assess the role of TRPV4 in HPKCs we used siRNA to knock down TRPV4 based on our previous experience (36). First, we demonstrated effective knockdown, namely a 70% reduction in *TRPV4* mRNA (qPCR) and 60% reduction in TRPV4 protein expression (Western blot) (SupFig.1A and B). siRNA-mediated knockdown of TRPV4 also significantly attenuated increase in pERK (SupFig.2A). In addition, siRNA-mediated knockdown of TRPV4 significantly reduced both IL-1ß and TNFα secretion from HPKC (Fig.2D, G). We also assessed the effect of compound-mediated inhibition of TRPV4 in these cells. TRPV4 inhibitor HC067047 significantly reduced IL-1ß and TNFα secretion in HPKCs (Fig.2E, H). Of note, IL-1ß is a direct inflammasome product whereas TNFα is not(38). We also tested additional TRPV4 specific inhibitors, and all significantly reduced IL-1ß secretion in HPCKs (SupFig.2B). Thus, secretion of the direct inflammasome product IL-1ß in HPKCs depends on KC TRPV4. This finding in a human primary KC-based experimental platform further corroborates that TRPV4 functions as a key calcium entry pathway upstream of the inflammasome in epidermal KC. To assess if TRPV4-dependent IL-1ß secretion was dependent on ERK signaling, we pretreated HPKCs with MEK-inhibitor U0126. Our findings were affirmative, namely U0126 significantly reduced IL-1ß secretion as a response to UVB, to similar levels as the siRNA *TRPV4* knock-down cells, suggesting ERK functions downstream of TRPV4 and upstream of IL1ß secretion. In keeping with this concept, U0126 did not significantly further inhibit IL-1ß secretion in the siRNA *TRPV4* knock-down cells (Fig. 2F), suggestive of both interventions affecting the same signaling pathway. In contrast, U0126 further reduced TNFα secretion in *TRPV4* siRNA mediated knockdown cells, suggesting that TNFα secretion, not a direct consequence of inflammasome activation, is only partially dependent on TRPV4 signaling (Fig.2I).

**Figure 2.**
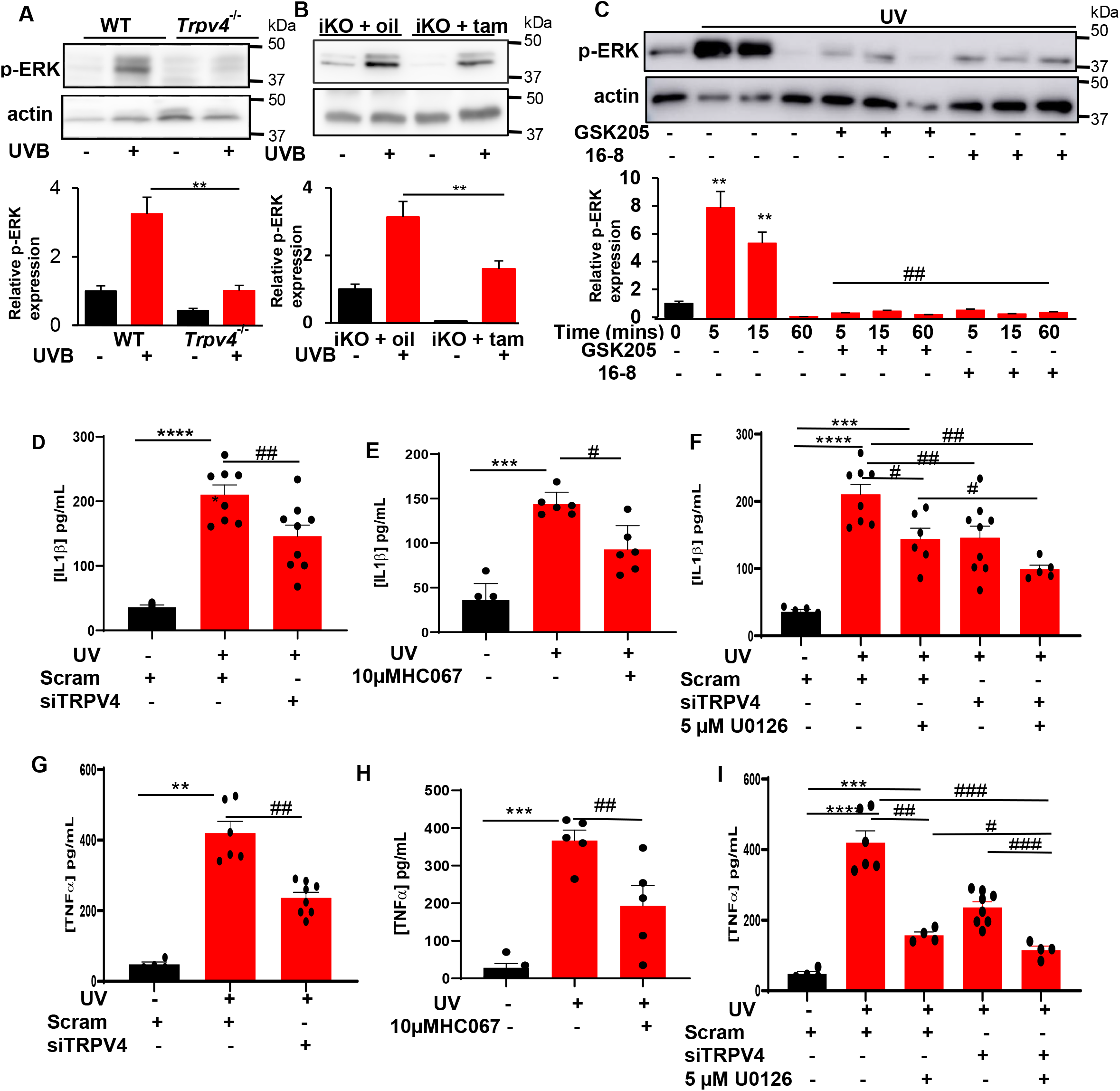
TRPV4 activation by UVB induces ERK phosphorylation, then triggers extracellular release of IL-1ß and TNFα in skin KCs. (A) ERK signaling is attenuated in back skin *Trpv4^−/−^* and (B) tamoxifen treated iKO animals. Western Blot analysis and quantification of bands demonstrates that UVB-mediated activation of KCs induces ERK activation and this is dependent on TRPV4. **p<0.01 vs. WT control. *n*=5 mice/group. (C) Time course of ERK activation in HPKCs after UV treatment. Western Blot analysis and quantification of bands demonstrates p-ERK is rapidly activated by 5 minutes after UV treatment, by 15 minutes p-ERK was 30% diminished and completely gone at 60 minutes. Pre-incubation withTRPV4 antagonists 10 μM GSK205 and 2 μM compound16-8 prevented the activation of ERK signaling at all the time points analyzed. **p<0.01 no UVB control and ^##^p<0.01 vs. UVB. *n* = 3 (D) IL1ß secretion is attenuated in HPKCs with siRNA mediated knockdown of TRPV4 and (E) pharmacological inhibition by pretreatment with TRPV4 antagonist 10 μM HC067047., ***p<0.001, ****p<0.0001 vs Veh. (DMSO). ^#^p<0.05 and ^##^p<0.01 vs. UVB. (F) MEK inhibitor U0126 (5 μM) inhibited UVB induced IL1ß secretion to similar levels as TRPV4 siRNA, and further reduced IL1ß secretion when combined with TRPV4 siRNA., ***p<0.001, ****p<0.0001, vs Veh. (DMSO). ^#^p<0.05 and ^##^p<0.01 vs. UVB. (G) siRNA mediated knockdown of TRPV4 and (H) pharmacological inhibition by pretreatment with TRPV4 antagonist 10 μM HC067047 inhibited TNFα. **p<0.01 and ***p<0.001 vs Veh. (DMSO). ^##^p<0.01 vs. UVB. (I) MEK inhibitor U0126 (5 μM) inhibited UVB induced TNFα secretion, and further reduced IL1ß secretion when combined with TRPV4 siRNA suggesting. ***p<0.001 and ****p<0.0001 vs. Veh. (medium) ^#^p<0.05 and ^##^p<0.01 and ^###^p<0.001 vs. UVB treatment. Two-tailed *t* test was used for statistical analysis. *n* = 5-8 independent cultures/experiment for HPKCs.

In HPKCs, it has been demonstrated previously that NLRP1 can function as the principal inflammasome sensor (32). UVB radiation induces NLRP1 activation in KCs, which is a key element of the skin’s photodermatitis/sunburn response (39–41). However other studies have also implicated the NLRP3 inflammasome (30,42). We therefore wanted to test the extent to which NLRP1 and NLRP3 were involved in UVB mediated proinflammatory secretory function in HPKCs. Using siRNA targeted to both NLRP1 and NLRP3 resulted in a 60% reduction at the mRNA level and 80% knockdown at the protein level when assessed 4 days after siRNA transfection (SupFig.3A, B). Indeed, siRNA mediated reduction in NLRP1 but not NLRP3 reduced IL-1ß secretion after UVB treatment (Fig.3A). To test a direct role of TRPV4 in the inflammasome pathway we stimulated HPKCs with TRPV4 selective activator, 4αPDD. As a result, we observed an increase in IL-1ß secretion (Fig.3B). This increase in IL-1ß was significantly inhibited in *NLRP1* siRNA knockdown HPKCs, to a lesser extent in *NLRP3* siRNA knockdown HPKCs. These findings reiterate the postulated signaling of TRPV4-mediated calcium influx upstream of KCs’ inflammasome, more specifically the NLRP1 inflammasome, for processing and secretion of proinflammatory and proalgesic IL-1ß. We also determined levels of TNFα, a pro-inflammatory/pro-algesic secreted mediator that is not a direct inflammasome product. We found that both NLRP1 and NLRP3 were involved in UVB and 4αPDD stimulated TNFα secretion (Fig. 3C-D). This TNFα-related result, together with the IL-1ß result indicate that TRPV4-mediated Ca^++^ influx directly activates the inflammasome which then leads to “classic” product of IL-1ß, yet also leads to increased secretion of TNFα as a more indirect effect - dependent on both NLRP1 and NLRP3 inflammasome.. Both findings cast in a new light the central and fundamental role of the inflammasome as a subcellular multiprotein machine that renders the keratinocyte a cellular generator of inflammation and pain. In this skin→neuron signaling mechanism, we note that TRPV4 functions upstream of the inflammasome in mammalian KCs.

**Figure 3.**
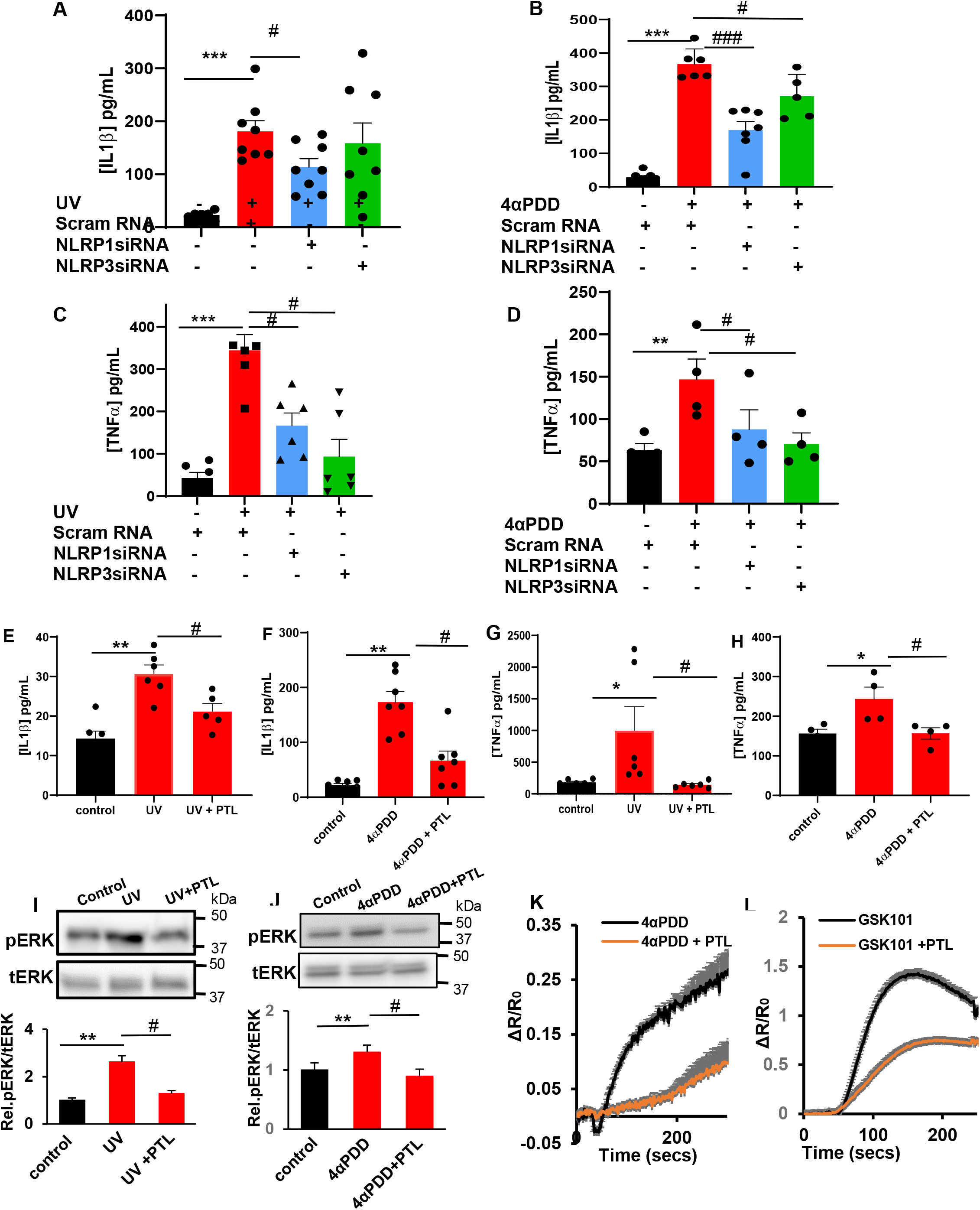
UVB and TRPV4 agonist 4αPDD activates NLRP1 inflammasome in HPKCs. KCs were transfected using siRNA targeted to NLRP1 and NLRP3. 24 hours after UVB and 4αPDD treatment of HPKCs, supernatants were analyzed by ELISA for IL1ß and TNFα secretion. (A) UVB induced IL1ß secretion is inhibited in siRNA mediated knockdown of NLRP1 but not siRNA knockdown of NLRP3 in HPKCs. The data are expressed as mean and SEM. ***p<0.001 vs. Veh. (medium) ^#^p<0.05 vs. UVB treatment. (B) TRPV4 agonist 4αPDD induced IL1ß secretion and this can be attenuated by siRNA mediated reduction of both NLRP1 and NLRP3. ***p<0.001 vs. Veh. (medium) ^#^p<0.05 and ^###^p<0.001 vs. UVB treatment. (C, D) TNFα secretion is mediated via the NLRP1 and NLRP3 inflammasome as siRNA mediated knockdown of NLRP1 and NLRP3 inhibited UVB and 4αPDD secretion of TNFα. **p<0.01 and ***p<0.001 vs. Veh. (medium) ^#^p<0.05 vs. UVB treatment. (E, F) Parthenolide (PTL) inhibits IL1ß secretion induced both by UVB and TRPV4 agonist 4αPDD. *p<0.05 and **p<0.01 vs. Veh. (medium) ^#^p<0.05 vs. UVB treatment. (G) UVB and TRPV4 agonist 4αPDD, H, induced TNFα secretion in inhibited by PTL. *p<0.05 vs. Veh. (medium) ^#^p<0.05 vs. UVB treatment. (I, J) Western Blot analysis and quantification of bands demonstrates PTL inhibits ERK phosphorylation at the 5-minute time point in both the UV and 4αPDD treated HPKCs *p<0.05 and **p<0.01 vs. Veh. (medium), #p<0.05 vs. UV/4αPDD treatment. (K, L) TRPV4 agonists, 4αPDD (50 μM) and GSK101 (20 nM) induced Ca^++^ influx into HPKCs and this influx is attenuated by pre-incubation with PTL (25 μM) n=200-300 cells/treatment. Two-tailed *t* test was used for statistical analysis. *n* = 4-7 independent cultures/experiment for HPKCs.

With translational relevance, to test how compound-mediated inhibition of the inflammasome relates to TRPV4 signaling in KC, we selected parthenolide (PTL), a sesquiterpene lactone from *Tanacetum parthenium* (feverfew), which is used as an herbal compound to treat inflammation and pain, including migraine headaches (43,44). Important for our approach, PTL is a known NLRP1 and NLRP3 inhibitor (45,46). PTL significantly inhibited both UVB and 4αPDD induced IL-1ß and TNFα secretion in HPKCs (Fig.3E-H). Given the rapid, TRPV4-dependent upregulation of pERK in response to UVB in these cells, we tested if PTL can inhibit pERK formation. We recorded affirmative results, namely that PTL inhibited ERK phosphorylation in both the UVB and 4αPDD-treated HPKCs (Fig.3I-J) suggesting PTL’s inhibitory action might affect signaling upstream of MEK/ERK. To inquire whether PTL can inhibit TRPV4 channel function, we studied Ca^++^ transients in HPKCs using chemical activation of TRPV4 with 4αPDD and GSK101. Our results show significant attenuation of the chemically-evoked Ca^++^ signal when pre-treating HPKCs with 20 μM PTL. Thus, the natural herbal compound PTL, which has a favorable profile in human clinical studies for migraine and other pain and inflammatory disorders, inhibits TRPV4 signaling in HPKCs. This finding confirms and extends our body of evidence presented here, suggesting TRPV4 functions upstream of the NLRP1 inflammasome in KCs, relevant for proinflammatory and proalgesic signaling in skin in response to UVB. How PTL inhibits TRPV4 at the level of ion channel functioning and how this finding can be translated preclinically will be the subject of future studies.

## DISCUSSION

Here we describe a previously unrecognized role for TRPV4 ion channels in skin KCs for permeating Ca^++^ ions that activate the inflammasome in response to UVB. Such a mechanism has long been sought after, and our results present concrete evidence in favor of this important process relying on TRPV4 channels, at least partially. We found that UVB triggers TRPV4 channel activation, which leads to Ca^++^-influx into KCs, which in turn activates the inflammasome machinery that leads to secretion of IL1ß, via direct inflammasome activation, and TNFα, via secondary effects.

TRP channels can be activated by cues such as heat, acidity, chemical activators, UVB, changes in osmolarity, and also by signals emanating from tissue injury such as ATP. Therefore, TRP channels have been postulated to be the origin of increased intracellular Ca^++^ in NLRP inflammasome activation. Here we communicate previously unreported findings that activation of TRPV4, either by UVB or selective TRPV4 activators, signals via the NLRP1 inflammasome in HPKCs. This particular argument rests on our observation that siRNA-mediated knockdown of NLRP1 significantly attenuated IL1ß secretion which is the result of activation of TRPV4 in HPKC.

In addition to TRPV4, KCs express other TRP channel such as TRPV1 and TRPV3, suggesting that these channels might also play an role in skin inflammation, as they control the secretion of IL1ß and other pro-inflammatory cytokines. Our current study suggests TRPV4 is an important initiator (evidenced by its role in the NLRP1 inflammasome pathway) and perpetuator (evidenced by its effects on TNFα) of inflammatory signaling in skin KCs in response to UVB.

In the future, it will be important to understand how TRPV4 activation leading to TRPV4-mediated Ca^++^-influx impacts interacting signaling machinery to result in inflammasome activation. Very likely, TRPV4 functions as a core ion channel ionophore within a multiprotein complex which can now be rationally discovered. It is already known that TRPV4 interacts with the cellular cytoskeleton, and this interaction is postulated to affect the functionality of TRPV4 channels as well as the cytoskeleton (47). We found that the natural compound parthenolide (PTL), known for its anti-inflammatory, analgesic and anti-tumor activity, has the capability to inhibit Ca^++^ influx via TRPV4 channels. These new insights suggest that PTL also acts upstream of the inflammasome. In regards to mechanisms of action of PTL affecting TRPV4 channels, this will now be a subject for rational discovery in future studies. Clearly, known effects of PTL on cytoskeletal functioning have to be considered as important potential contributors. Intriguingly, PTL has been shown to affect detyrosination of tubulin leading to its stabilization (48,49). As an interesting complement to our observations, PTL inhibited TRPV4-dependent Ca^++^-influx caused by fluid sheer stress in osteocyte mechanotransduction (48,49). In our experiments reported here, pretreatment with PTL may have enhanced microtubule stiffness subsequently reducing TRPV4’s ability to permeate Ca^++^, which in turn reduced downstream inflammasome activation. This would indicate that microtubule detyrosination and microtubules’ subsequent altered mechanical properties, namely increased stiffness, can also potently regulate TRPV4 function, yet in the absence of a mechanical stimulus.

The translational relevance of our findings is rooted in the fact that we now have an increased incentive to target TRPV4 for treatment of pain and tissue damage associated with sunburn and other NLRP1-related skin diseases such skin inflammation and epidermal hyperplasias (50). PTL is currently taken orally as a treatment for migraine headaches, fever, oral, odontogenic and arthritis pain, with some recent premise in oncology to contain malignant growth and tumor spread. Its use can be expanded toward dermatologic disorders associated with skin inflammasome activation. Furthermore, PTL and TRPV4-selective inhibitors can be formulated for topical application to epidermis in such disorders, in particular UVB overexposure-induced pathological conditions, possibly also skin cancers (51).

Another interesting possibility emerges in melanoma, a deadly skin cancer where TRPV4 activation can be considered a translational goal. In melanoma cancer cells, activation of TRPV4 has been shown to enhance cell apoptosis (24,52). The experiments suggested that TRPV4 channel activation contributed to melanoma cell death via Ca^++^-influx, which also activated the AKT pathway with net anti-proliferative effects (52). One possible translational application is to enhance melanoma cell death via use of targeted UVB and/or topical TRPV4 agonists in conjunction with currently established melanoma treatments.

## MATERIAL AND METHODS

### Animals

Wild-type C57bl/6j mice were purchased from the Jackson Laboratory*. Trpv4* knockout (KO) mice were generated in our laboratory as previously described(53). Keratinocyte-specific, tamoxifen (tam)-inducible *Trpv4* knockdown mice were used as previously described(23).Using the same genomic mouse *Trpv4* clone as reported for the *Trpv4^−/−^* pan-null mouse and the *Trpv4* locus was engineered by insertion of loxP sites surrounding exon 13 which encodes transmembrane domains 5-6 This mutation was propagated in mice that were crossed to K14-Cre-ER^tam^ mice, so that K14-Cre-ER^tam^::*Trpv4*^lox/lox^ mice could be induced by tamoxifen (tam) administration via oral gavage for five consecutive days at 5mg/day in 0.25 ml corn oil at 2–2.5 months of age, plus a booster 2 weeks after the last application. Control animals received the same volume of corn oil. Efficiency of targeting was verified by quantitative real-time PCR and immunohistochemistry for *Trpv4* expression in skin at gene and protein levels, respectively (23).

Mice were housed in climate-controlled rooms on a 12/12-h light/dark cycle with water and a standardized rodent diet available *ad libitum*. All animal protocols were approved by the Duke University Institutional Animal Care and Use Committee (IACUC) in compliance with National Institutes of Health (NIH) guidelines. All of these mouse lines have C57bl/6 background and were PCR-genotyped before use.

### Chemicals and Antibodies

We used the following compounds: 4α-phorbol 12,13 didecanoate (4α-PDD; TRPV4 activator; Tocris), GSK1016790A, HC067047, and U0126 were purchased from Sigma (St. Louis, MO). RN-1734 (TRPV4 inhibitor) was purchased from Tocris, Parthenolide was purchased from Cayman Chemicals. GSK205, pan TRPV4 blocker was synthesized by the Small Molecule Synthesis Facility at Duke University (37). Rabbit monoclonal anti-ERK and polyclonal anti-phospho-MEK were obtained from Cell Signaling Technology (Danvers, MA), polyclonal anti-TRPV4 from Abcam (Cambridge, MA), and polyclonal anti-actin from Sigma. 4′,6-diamidino-2-phenylindole (DAPI) was obtained from Sigma.

### Cell Culture and Transfection

Primary mouse keratinocytes were derived from back skin of P2-P4 newborn mice as previously described(53). The epidermis was separated from the dermis by a 1-hour dispase (BD Biosciences, MA) treatment and then the KCs were dissociated from the epidermis using trypsin. KCs were plated on collagen coated dishes and grown in keratinocyte serum free media (Gibco) supplemented with bovine pituitary extract and epidermal growth factor (EGF) (R&D Systems, Minneapolis, MN, USA), 10^−10^ molL-^1^ cholera toxin (Calbiochem, San Diego, CA, USA) and 1 X antibiotics/antimycotics (Gibco), in an incubator at 5% CO_2_ and 37°C.

Human primary keratinocytes (HPKCs) were cultured as previously described (13). In brief, surgically discarded foreskin samples, obtained from Duke Children’s Hospital in accordance to institutionally approved IRB protocol, were incubated with Dispase (Gibco, 4 U/ml) for 12-16 h at 4°C followed by 0.05% trypsin (Gibco) for 10-20 min at 37°C. Cells were maintained in keratinocyte serum-free media (Invitrogen) with 5% CO_2_ at 37°C and used at passage 2-3.

HPKCs were transfected with presdesigned siRNA from using either HiPerfect transfection reagent (Qiagen) or Lipfectamine RNAiMax (Invitrogen) according to manufacturer instructions. Silencer Select Pre-designed siRNA were purchased from Ambion, Negative control #1 siRNA (Cat#430843), NLRP3 (Cat #4392420 ID s534395, NLRP1 (Cat #4392420 ID:s22522), TRPV1 (Cat #4392420 ID:s14818), TRPV3 (Cat #4392420 ID:s46346), TRPV4 (Cat #4392420 ID:s34003). Taqman Gene Expression Assays containing predesigned primers were purchased from Applied Biosystems for qPCR to determine efficiency of knockdown. (Hs00248187_m1NLRP1, Hs00918082_m1NLRP3, Hs01049631_NLRP1 FAM, Hs00218912_m1 TRPV1, Hs00376854_m1 TRPV3, Hs00540967_m1 TRPV4, Hs00354836_m1 CASP1, Hs99999905_m1 GAPDH, Hs01556773_m1 EIF4A3. Knockdown efficiency was also detected by western blot analysis.

### Intracellular Calcium Imaging

Ca^++^ imaging of mouse primary KCs cells was conducted using 2 μM Fura-2 acetoxymethyl ester for loading and following a protocol for ratiometric Ca^++^ imaging using 340/380 nm blue light for dual excitation, recording emissions with specific filter sets. Ratios of the emissions were acquired every 3 sec. ΔR/R_0_ is the fraction of the increase of a given ratio over the baseline ratio, divided by baseline ratio.

### Keratinocyte UV Irradiation

The 1°MKCs and HPKCs were grown in KSFM (supplemented with EGF and BPE) and medium was exchanged with fresh KSFM before experiment. HPKCs were pre-incubated with inhibitors or vehicle for 15 mins, then irradiated with 60 mJ/cm^2^ UVB using Spectroline Medium wave UV 312nm lamp or left untreated for controls. Twenty-four hours later the supernatant were collected for ELISA. ELISA (Human IL1 beta/IF-F2 DuoSet ELISA (DY201), and Mouse IL1 beta/IF-F2 DuoSet ELISA(DY401) from R&D Sytems, and Human TNF-alpha Stand TMB ELISA (900-T25) was from PeproTech. Lysates were collected for either western blotting or qPCR.

### Mice hind paw and back skin UV Irradiation

Mice were confined by Plexiglas enclosures on top of a 25-× 26-cm Bio-Rad Gel Doc 2000 UV transilluminator (302 nm), and otherwise allowed to openly explore this environment. UV exposure lasted for 5 min with an exposure of 600 mJ/cm^2^ (6), equivalent to 5–10 minimal erythema-inducing dose (MED). Thorough observations upon initiation of this method demonstrated that hind paws were exposed to UV throughout this period and that animals did not engage in licking behavior during UVB exposure. Mice dorsal back skin was shaved and were irradiated under general anesthesia using the Spectroline Medium wave UV 312nm lamp for 5 min with an exposure of 600 mJ/cm^2^ (6), 15 minutes after initial UV exposure, animals were sacrificed and the irradiated regions of back skin were collected for western blotting to assess levels of pERK.

### GSK205 topical treatment

A viscous solution of 68% (vol/vol) EtOH/5% glycerol plus 1 mM or 5 mM GSK205 (none for control) was applied to hind paws by rubbing in 20 μL, applied at time points 1 h and again 10 min before UV exposure.

### Tissue Processing and Immunohistochemistry

Mice were perfused intracardially with 30 ml phosphate buffered saline, pH 7.4 (PBS), followed by 30 ml solution of 4% paraformaldehyde in PBS. Tissues were dissected out, post-fixed in 4% paraformaldehyde. Tissue blocks were further cryoprotected in 30% sucrose in PBS for 24–48 h and sectioned on a cryostat. Sections of footpads (10 μm) were thaw-mounted onto slides. Sections were blocked with 5% normal goat serum (NGS; Jackson) in PBS/0.05% Tween20 (PBS-T), and incubated overnight with primary antibodies, rabbit anti-TRPV4 (1:300; Abcam), goat anti-IL1ß, (1:200; Santa Cruz Biotechnology Inc); rabbit anti-caspase-1(1:200; Biovision Research Products, CA). After washing, sections were incubated with secondary antibodies (AlexaFluor595 and AlexaFluor488-conjugated antibodies at 1:600; Invitrogen) for 2 h, rinsed, mounted, and cover-slipped with Flouromount (Sigma). Digital micrographs were obtained using a BX60 Olympus upright microscope equipped with high-res CCD camera and acquired with constant acquisition and exposure settings using ISEE software. Images were analyzed using imageJ open source software.

## STATISTICAL ANALYSIS

All data are expressed as the mean ± S.E. Two-tailed *t* tests or one-way analysis of variance followed by Dunnett post hoc test were used for group comparisons. *p* < 0.05 indicated statistically significant differences.

## ACKNOWLEDGMENT

“Research reported in this publication was supported by the National Center for Advancing Translational Sciences of the National Institutes of Health under Award Number 1KL2TR002554. The content is solely the responsibility of the authors and does not necessarily represent the official views of the National Institutes of Health.”

## SUPPORTING INFORMATION

**Supplementary Figure 1.**
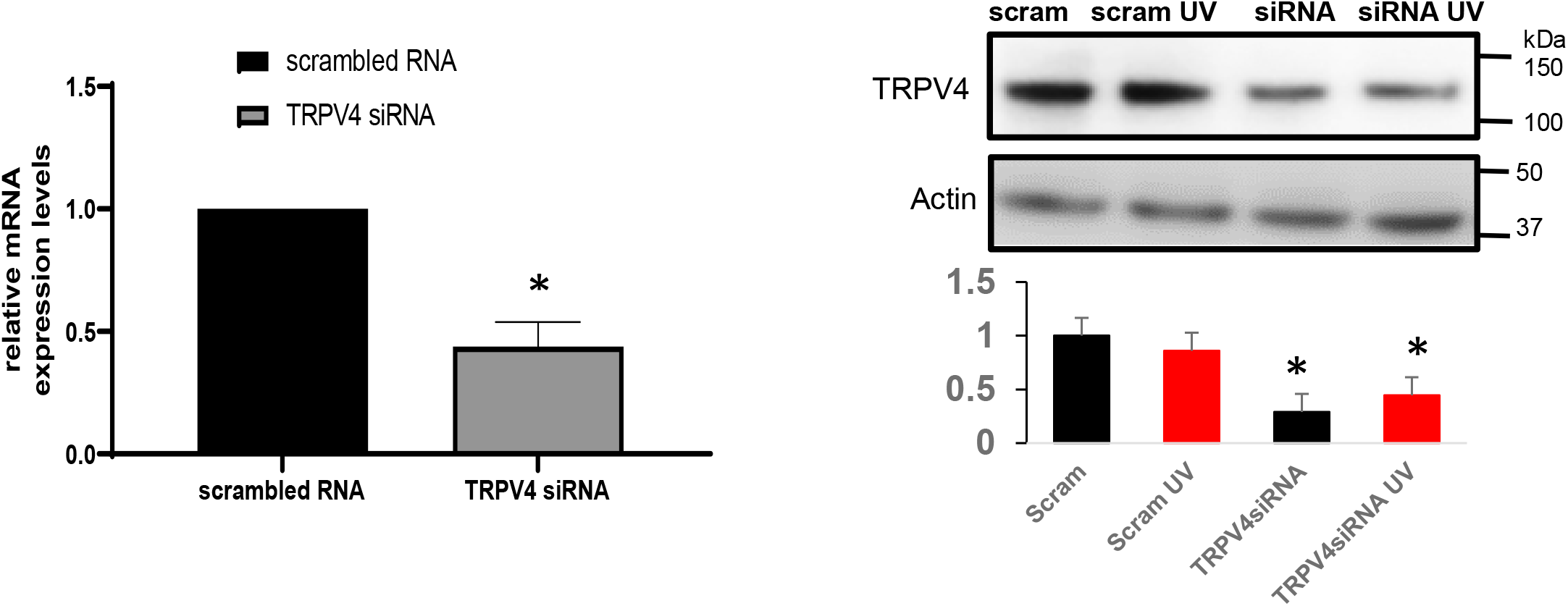
A. Relative mRNA level of TRPV4 showing approximately 60% knockdown with siRNA to TRPV4 in HPKCs. B. Western blot showing TRPV4 protein expression levels normalized to actin showing approximately 70% knockdown of TRPV4 using the siRNA targeted to TRPV4. Two-tailed *t* test was used for statistical analysis. *n* = 3 independent cultures/experiment for HPKCs.

**Supplementary Figure 2.**
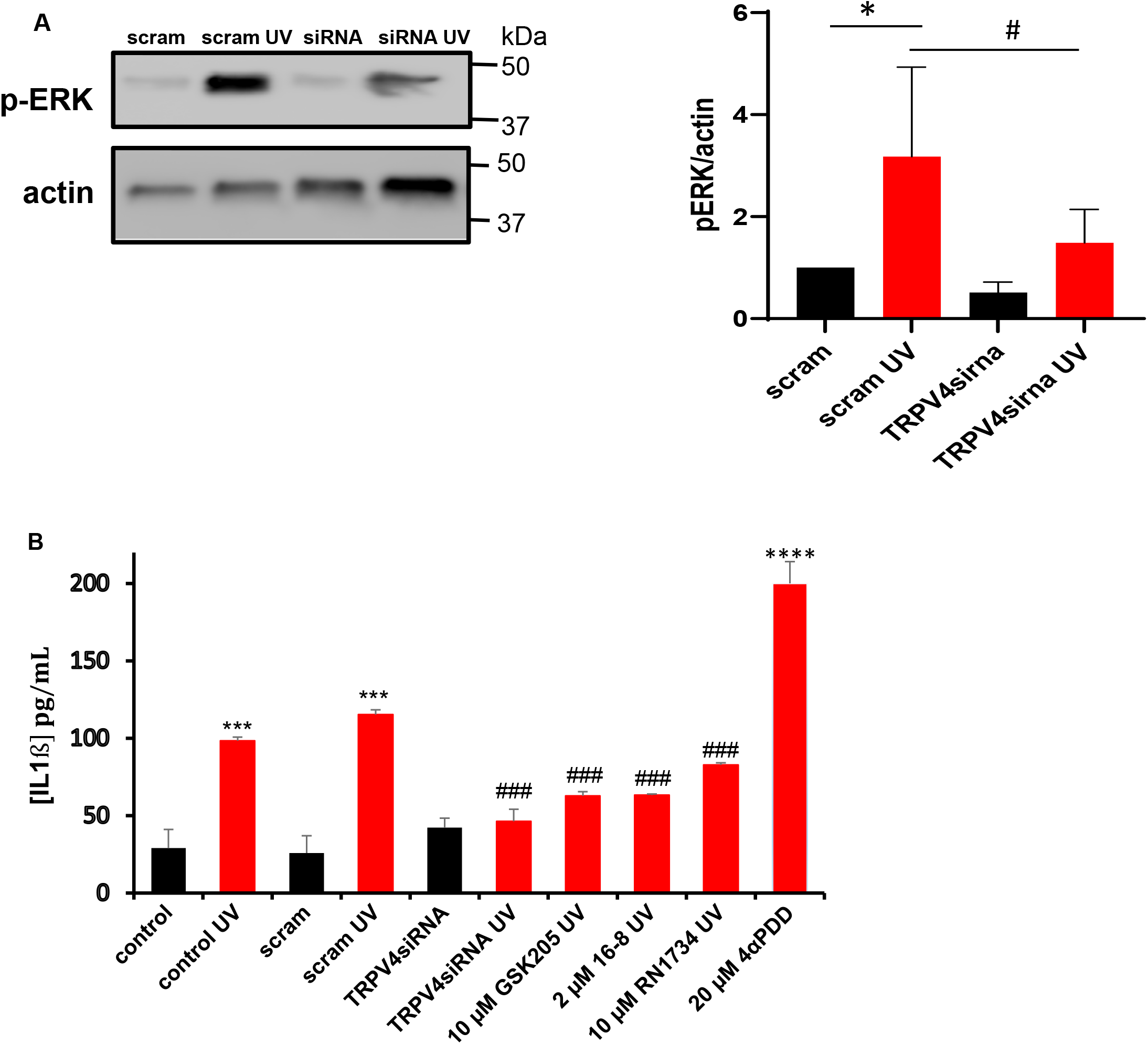
ERK signaling is attenuated in siRNA mediated knockdown of TRPV4 in HPKCs after 5 minutes after UV treatment, compared to the rapid p-ERK activation in the scrambled RNA treated control HPKCs which is blocked in the absence of each TRP channel. B. siRNA mediated knockdown of TRPV4 and pharmacological inhibition by pretreatment with TRPV4 antagonists 10 μM GSK205, 10 μM RN1734, and 2 μM compound 16-8 abrogates IL1ß secretion in HPKCs. Treatment with 20 μM of 4αPDD, a TRPV4 agonist is a potent inducer of IL1ß secretion. **p<0.001, and ****p<0.0001vs Veh. (DMSO). ###p<0.001 vs. UVB Two-tailed *t* test was used for statistical analysis. *n* = 3 independent cultures/experiment for HPKCs.

**Supplementary Figure 3.**
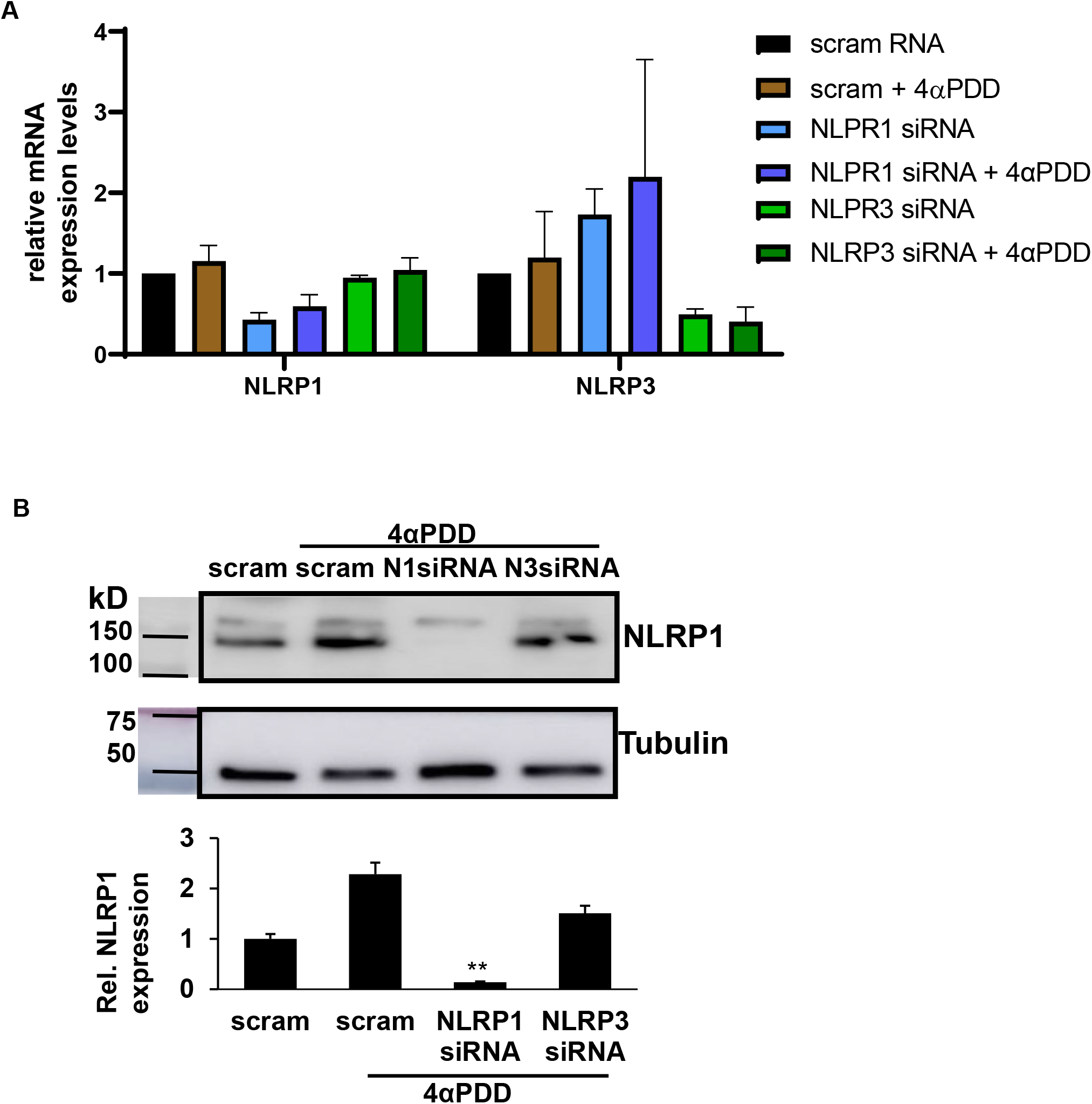
HPKCs were transfected using siRNA targeted to NLRP1(N1) and NLRP3(N3). 7hours after transfection cells were treated with 20μM 4αPDD and 24 hours after treatment, cell lysates were collected and analyzed via western blot and qPCR. A. Relative mRNA level using siRNA to NLRP1 and NLRP3 showing approximately 60% knockdown. B. TRPV4 agonist 4αPDD activates NLRP1 inflammasome in HPKCs. 4αPDD enhanced NLRP1 protein expression which was absent in siRNA transfected cells. Treatment with 20 μM of 4αPDD, a TRPV4 agonist is a potent inducer of IL1ß secretion. **p<0.001, and ****p<0.0001vs Veh. (DMSO). ###p<0.001 vs. UVB Two-tailed *t* test was used for statistical analysis. *n* = 3 independent cultures/experiment for HPKCs.

## Notes

### Competing Interest Statement

The authors have declared no competing interest.

### Summary of Updates

Minor edits to figures, write-up of results and discussions, editing the referencing, adapting the style of the paper for submission to a selected target journal.

